# Screening Short Peptide Binders to IL-17A Using Classical Molecular Dynamics

**DOI:** 10.64898/2026.09.04.749315

**Authors:** Xu Wu, Wei Fu

## Abstract

Identifying short peptides that bind therapeutically relevant proteins is a practical step toward developing peptide-based inhibitors. Here we screened 64 tetrapeptides for binding to the IL-17A homodimer using classical all-atom molecular dynamics simulations (GROMACS 2022.2, FF19SB force field, TIP3P water, 200 ns production per complex). Binding was ranked by a composite score that combines short-range interaction energies (Coul-SR + LJ-SR), close-contact frequency, mean intermolecular distance, and hydrogen bond occupancy. The tetrapeptide LYS-THR-CYS-ASP achieved the highest score in the set (composite score 2.85, total interaction energy −556 kJ/mol, average 8.0 H-bonds). Across all 64 simulations, electrostatic interactions accounted for approximately 72% of the total interaction energy. Stronger binders tended to contain charged and polar residues (Asp, Glu, Cys, Arg, Thr), while weaker binders were enriched in hydrophobic residues. Contact analysis identified a recurring interaction surface on IL-17A centered on cysteine and aromatic residues (Chain A: CYS87, Chain B:CYS196, A:HIS86, B:TYR175, B:HIS195), which we describe as a cysteine-aromatic cradle. These results nominate LYS-THR-CYS-ASP as a candidate for experimental binding validation and suggest simple sequence guidelines for designing IL-17A-targeting peptides.

## 1. Introduction

IL-17A is a pro-inflammatory cytokine that plays a central role in chronic inflammatory diseases including psoriasis, psoriatic arthritis, rheumatoid arthritis, and ankylosing spondylitis [1]. Monoclonal antibodies targeting IL-17A (secukinumab, ixekizumab) and IL-17A/F (bimekizumab) are clinically effective, with combined annual sales exceeding $7 billion [2]. However, antibodies are expensive to manufacture, require cold-chain storage and parenteral administration, and can elicit anti-drug antibodies [3]. These drawbacks motivate the search for smaller antagonists that could be developed into orally available or topically delivered drugs.

Small-molecule and peptide-based approaches to targeting the IL-17A/IL-17RA interface have been reviewed by Zhang and Dömling, who documented several chemotypes in preclinical and early clinical development [4]. Liu et al. identified a 15-residue peptide antagonist (HAP) through phage display that binds the IL-17A dimer and competes with IL-17RA [5], and Kochhar et al. subsequently used computational mutagenesis to propose improved variants [6]. These studies establish that the IL-17A dimer surface can accommodate peptide ligands, but they do not systematically examine how peptide sequence determines binding affinity across a diverse library.

Tetrapeptides are a practical system for such a systematic study. The sequence space (20^4^ = 160,000) is large enough to be chemically diverse yet small enough to be sampled meaningfully by computation [7]. Wang and colleagues have used coarse-grained MD and deep learning to screen the full tetrapeptide space for aggregation propensity, finding that aromatic and hydrophobic residues drive self-assembly while charged residues suppress it [8,9]. A related co-modeling framework combining sequence and chemical graph representations improved prediction accuracy for peptide properties [10].

In this study, we apply classical all-atom MD simulations to screen 64 tetrapeptides selected from a 1,000-member library for binding to IL-17A. Each tetrapeptide–IL-17A complex was simulated for 200 ns. Binding was assessed through a multi-metric score incorporating interaction energies, contact frequencies, intermolecular distances, and hydrogen bond counts. We identify a top candidate, characterize the IL-17A residues most frequently contacted by strong binders, and derive simple sequence preferences that may guide future peptide design.

## 2. Methods

### 2.1 System Preparation

The IL-17A structure was obtained from AlphaFold3 as a homodimer of chains A and B (218 residues each). IL-17A belongs to the cysteine knot fold family, with intrachain disulfide bonds (Cys71-Cys121, Cys76-Cys123) and an interchain disulfide [12,13]. From a 1,000-member tetrapeptide library, 64 sequences were selected to cover diverse amino acid compositions. Each tetrapeptide was placed near IL-17A using gmx insert-molecules in a 7.0 × 7.0 × 7.0 nm^3^ cubic box, solvated with TIP3P water, and neutralized with 150 mM Na^+^/Cl^−^. Simulations used the FF19SB all-atom force field [14].

### 2.2 Simulation Protocol

All simulations were performed with GROMACS 2022.2 [15] using Intel oneAPI 2022.1. The protocol consisted of: (1) steepest-descent energy minimization, (2) NVT equilibration (100 ps, 300 K, V-rescale thermostat), (3) NPT equilibration (100 ps, 1 bar, Parrinello-Rahman barostat), (4) NVT production (100 ns), (5) second NVT equilibration (100 ps), and (6) NPT production (200 ns). The 200 ns trajectory (coordinates saved every 20 ps, 10,001 frames) was used for analysis. A 2 fs timestep was employed with LINCS constraints on hydrogen bonds. Short-range nonbonded interactions were cut off at 1.0 nm; long-range electrostatics used Particle Mesh Ewald (0.16 nm grid spacing). Dispersion correction was applied to van der Waals interactions. Each system ran on 32 MPI processes on AMD EPYC 64-core nodes.

### 2.3 Interaction Energy Decomposition

Interaction energies between IL-17A and the tetrapeptide were computed using gmx mdrun -rerun with energy group decomposition (energygrps = IL17 Peptide). Short-range Coulomb (Coul-SR) and Lennard-Jones (LJ-SR) terms were extracted using gmx energy and averaged over 10,001 frames. The total interaction energy was taken as Etotal = ECoul-SR + ELJ-SR.

### 2.4 Contact, Hydrogen Bond, and Structural Analysis

Per-residue minimum distances were computed with gmx mindist -res. Residue pairs with distances below 0.5 nm were classified as contacting; those below 0.3 nm as close contacts. Intermolecular hydrogen bonds were identified with gmx hbond using default criteria (donor-acceptor distance ≤ 0.35 nm, angle ≤ 30°) [16]. Backbone RMSD values for the peptide and IL-17A were computed with gmx rms relative to the starting structures.

### 2.5 Composite Binding Score

Peptides were ranked using a composite score combining four normalized quantities: Binding Score = 0.30 × S_contacts_ + 0.20 × S_distance_ + 0.20 × S_hbond_ + 0.30 × S_energy_. Each term was normalized to [0, 1]. S_contacts_ reflects close contacts (<0.3 nm) normalized to a ceiling of 200. S_distance_ uses the mean intermolecular distance dmean: max (0, 1 − (d_mean_ − 0.15)/0.85). S_hbond_ normalizes the average H-bond count to a ceiling of 8. S_energy_ normalizes the interaction energy magnitude to a ceiling of |−600| kJ/mol. The score was scaled to a nominal 0-4 range.

### 2.6 Enrichment Analysis and Hotspot Mapping

Amino acid enrichment was calculated as R = (N_top_ + 0.5)/(N_bottom_ + 0.5) comparing the 40 residue positions from the 10 highest-scoring peptides against the 40 positions from the 10 lowest-scoring peptides. IL-17A hotspot residues were identified by per-residue contact frequency across the top 20 peptides, with a contact defined as minimum distance < 0.5 nm. Residues contacted by ≥12 of 20 top peptides were designated as consensus hotspots [17].

## 3 Results

### 3.1 Binding Score Ranking

Composite binding scores for the 64 tetrapeptides ranged from 1.31 to 2.85 (mean 1.74 ± 0.26, Figure 1). LYS-THR-CYS-ASP achieved the highest score of 2.85, followed by ARG-TRP-THR-ARG (2.10), PHE-SER-PRO-TYR (2.10), and ASN-HIS-GLU-GLN (2.07). The top-ranked peptide recorded the most favorable total interaction energy (−556 kJ/mol), the largest Coulombic contribution (−510 kJ/mol), and the highest average number of hydrogen bonds (8.0 per frame, Table 1).

**Table 1.**
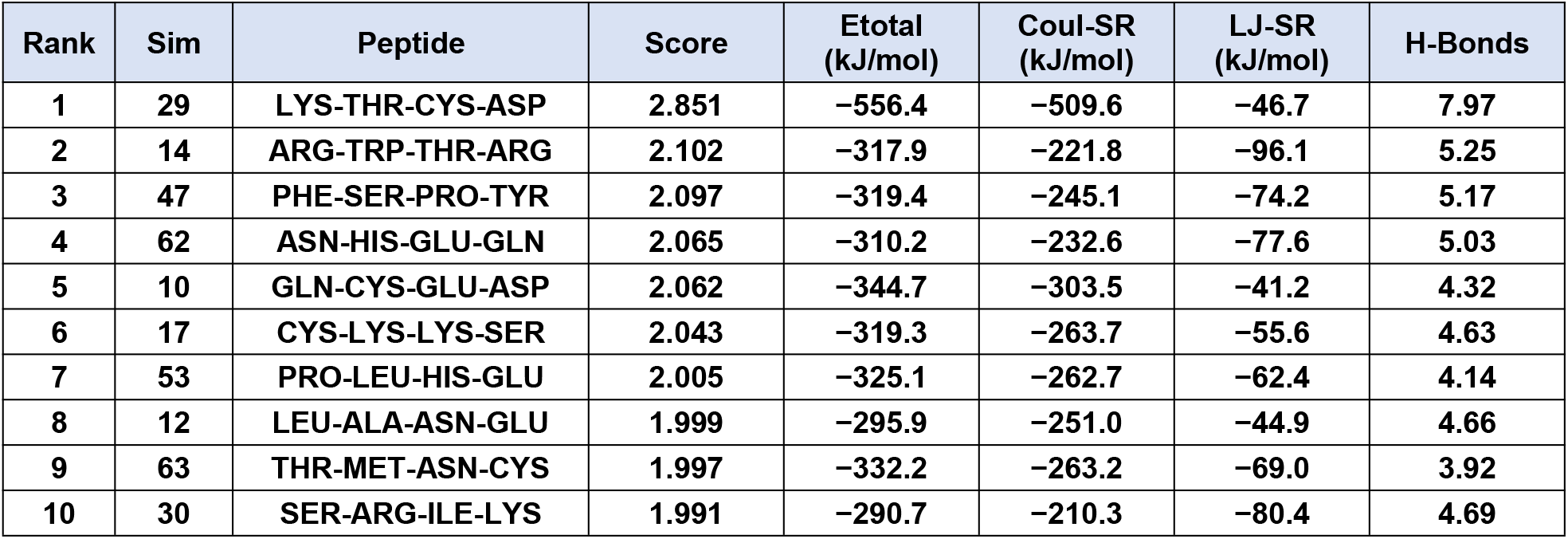
Top 10 tetrapeptides ranked by composite binding score.

**Figure 1.**
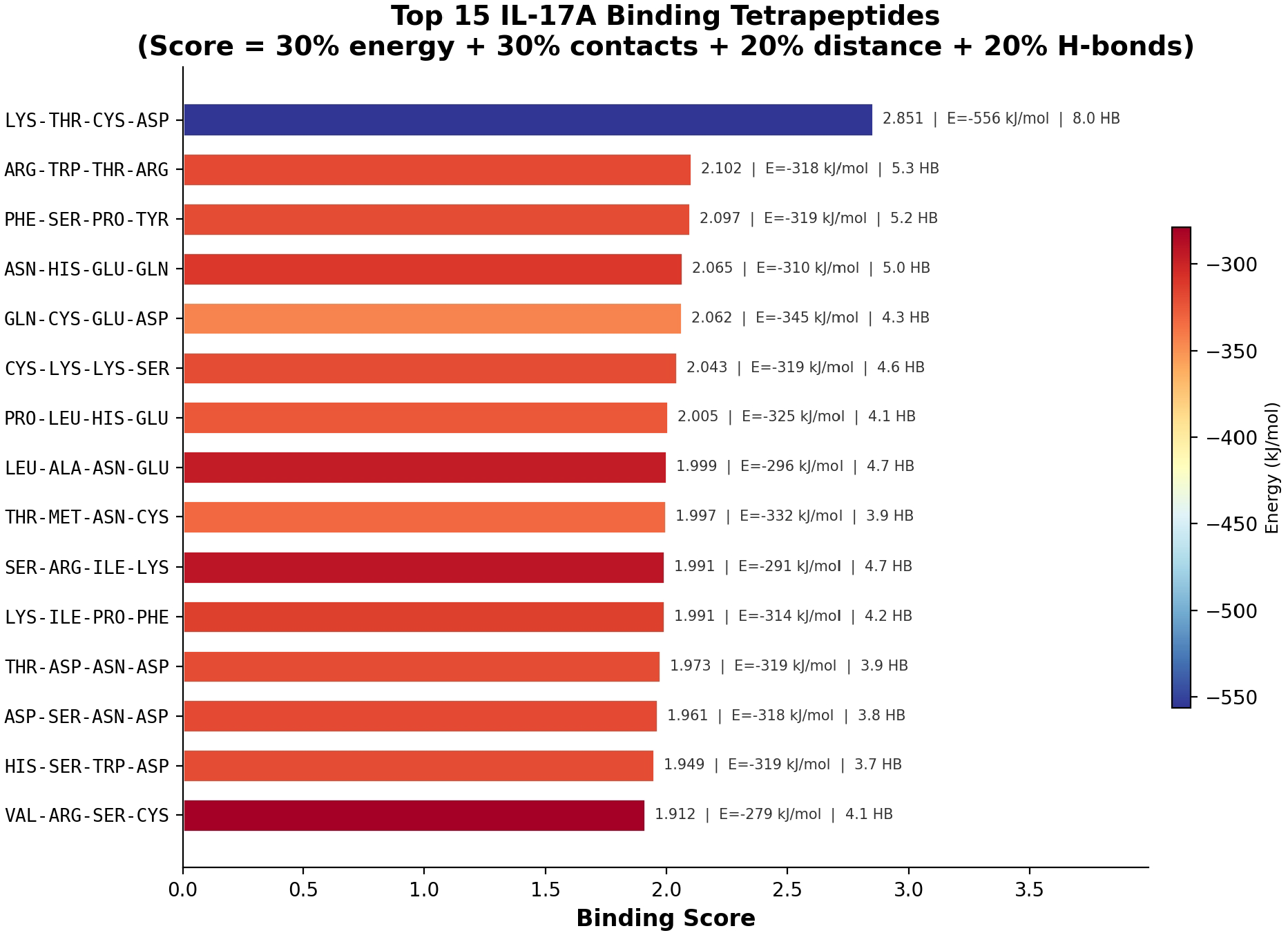
Top 15 tetrapeptides ranked by composite binding score. Bars are colored by total interaction energy. The dashed line marks the dataset mean (1.74).

### 3.2 Interaction Energy Analysis

Across all 64 simulations, short-range electrostatic interactions (Coul-SR) contributed approximately 72% of the total interaction energy (−179 ± 74 kJ/mol for Coul-SR vs. −68 ± 21 kJ/mol for LJ-SR), giving a Coulomb-to-LJ ratio of roughly 2.6:1 (Figure 2). This trend was consistent across the dataset: the five highest-scoring peptides all fell in the upper range of the Coul-SR distribution, while their LJ-SR values were unremarkable. For example, the top-ranked LYS-THR-CYS-ASP had an LJ-SR of −47 kJ/mol, which is below the dataset average of −68 kJ/mol. Hydrogen bond count and Coul-SR were correlated (Pearson r = 0.72), suggesting that the H-bond network is the main structural route through which favorable electrostatic contacts are made at this interface [16].

**Figure 2.**
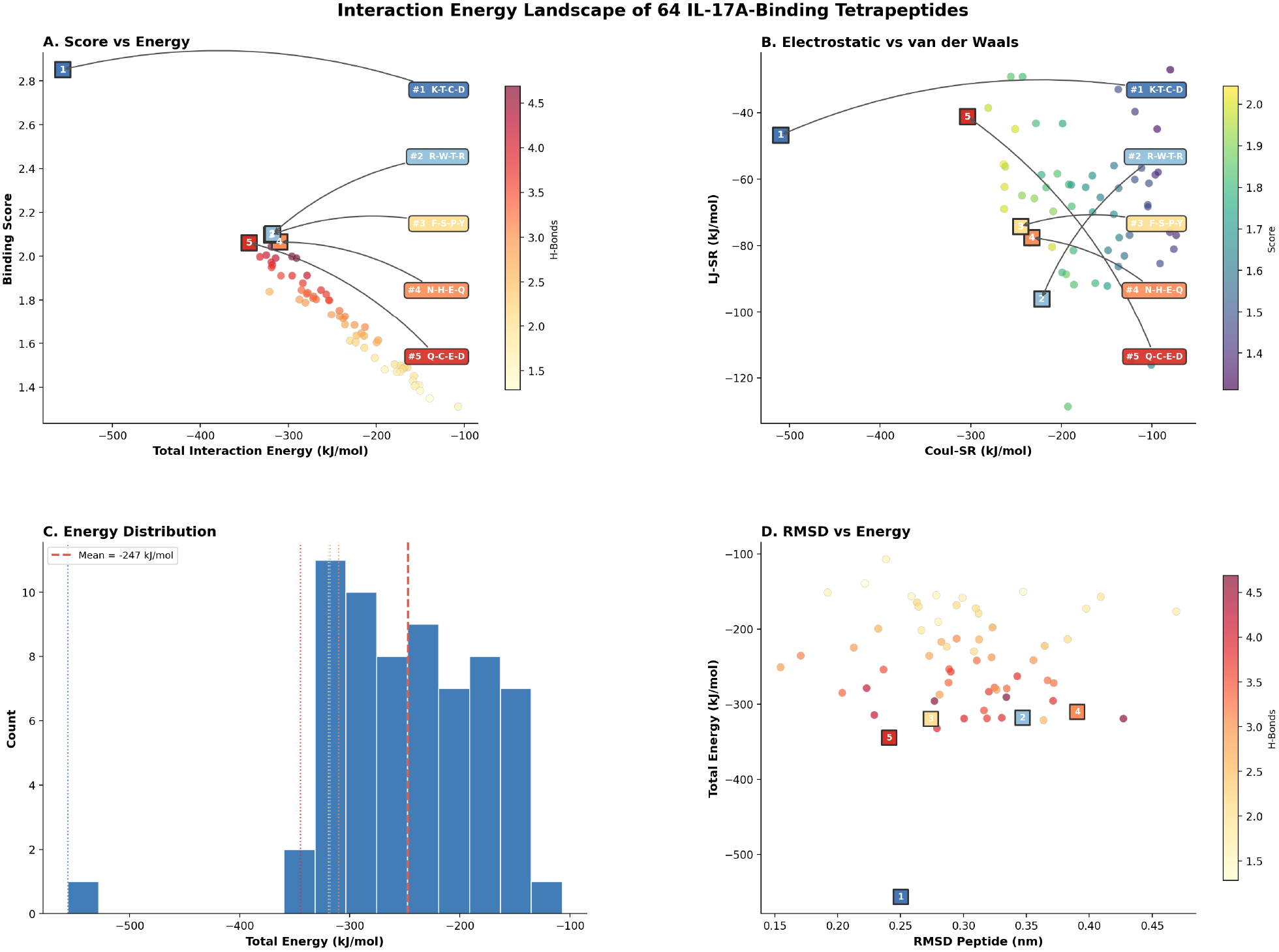
Interaction energy analysis. (A) Binding score vs. total interaction energy, colored by H-bond count. Top 5 peptides are marked. (B) Coul-SR vs. LJ-SR, colored by binding score. (C) Distribution of total interaction energies (dashed line: mean). (D) Peptide backbone RMSD vs. total interaction energy.

Per-residue contact analysis across the top 20 peptides identified 14 IL-17A residues contacted by 12 or more of the top binders (Figure 3, Table 2). Three residues were contacted by 15 of 20 (75%): A:ARG1, A:CYS87, and B:TYR175. Both chains contributed roughly equally to the interface (Chain A: 106 residues; Chain B: 108 residues).

**Table 2.** IL-17A residues contacted by ≥12 of 20 top-binding peptides.

| Residue | Chain | Peptides Contacting | Best Dist. (nm) |
| --- | --- | --- | --- |
| ARG1 | A | 15/20 (75%) | 0.146 |
| CYS87 | A | 15/20 (75%) | 0.179 |
| TYR175 | B | 15/20 (75%) | 0.150 |
| PRO0 | A | 14/20 (70%) | 0.150 |
| MET4 | A | 14/20 (70%) | 0.181 |
| HIS86 | A | 14/20 (70%) | 0.164 |
| MET113 | B | 14/20 (70%) | 0.171 |
| CYS196 | B | 14/20 (70%) | 0.162 |
| PRO197 | B | 14/20 (70%) | 0.153 |
| PRO84 | A | 13/20 (65%) | 0.167 |
| ASN89 | A | 13/20 (65%) | 0.150 |
| ASP170 | B | 13/20 (65%) | 0.142 |
| HIS195 | B | 13/20 (65%) | 0.152 |
| ASN198 | B | 13/20 (65%) | 0.162 |

**Figure 3.**
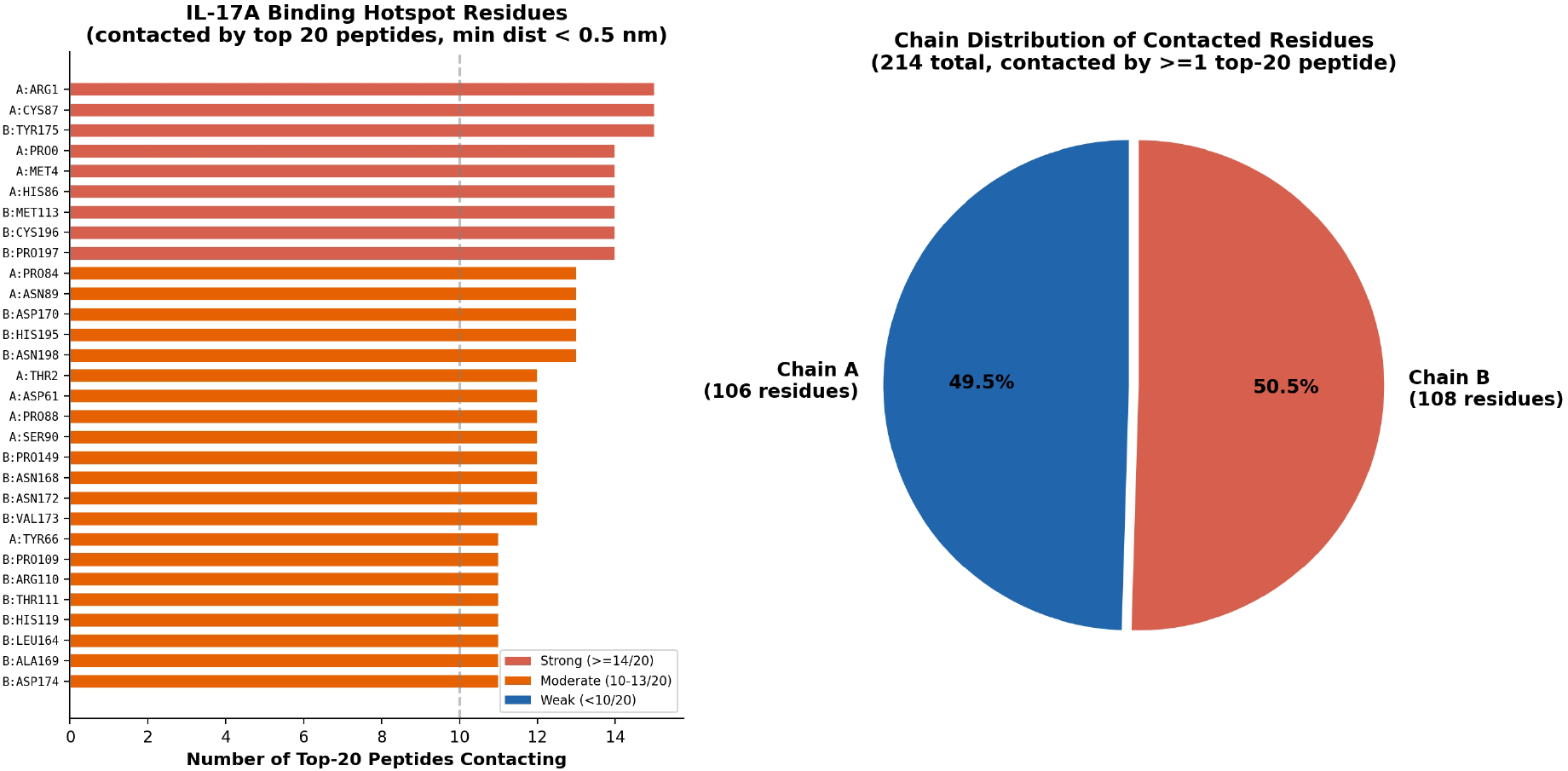
IL-17A contact hotspots. (A) Per-residue contact frequency among the top 20 peptides (min. distance < 0.5 nm). Red: ≥14/20, orange: 10–13/20, blue: <10/20. (B) Chain distribution of contacted residues.

**Figure 4.**
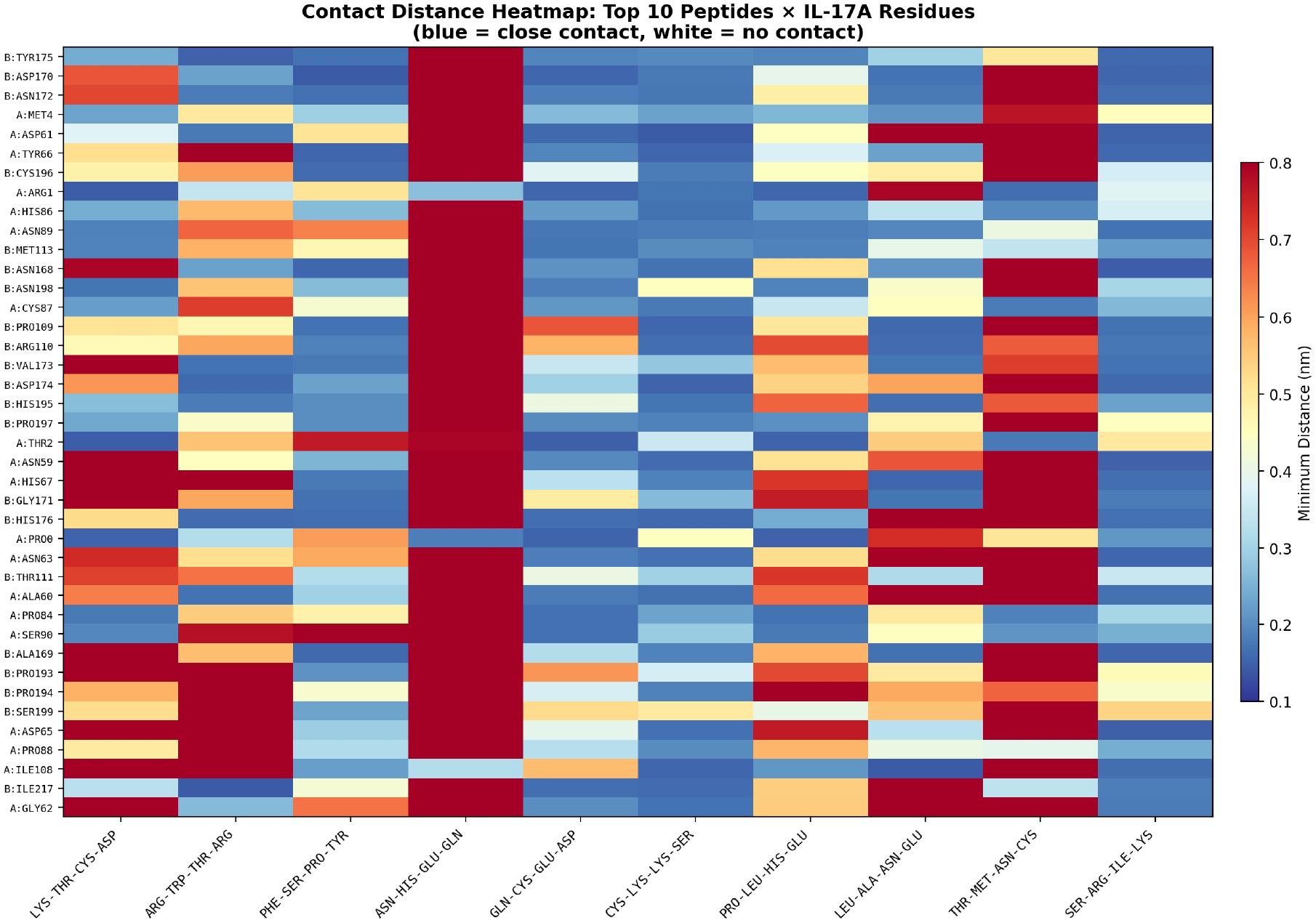
Contact distance heatmap for the top 10 peptides against the 40 most-contacted IL-17A residues. Blue: close contact (<0.2 nm); red/pale: distant or no contact.

The hotspot region is characterized by two cysteine residues (A:CYS87, B:CYS196) flanked by histidines (A:HIS86, B:HIS195) and a tyrosine (B:TYR175), set within a proline-rich scaffold (PRO0, PRO84, PRO197). We refer to this as a cysteine-aromatic cradle: a compact (∼1.5 nm^2^) surface patch that exposes polarizable sulfur atoms, π systems, and titratable groups. The electrostatic character of this patch, enriched in arginine, histidine, and cysteine, likely favors negatively charged and polarizable side chains on binding peptides [17].

### 3.3 IL-17A Contact Hotspots

### 3.4 Amino Acid Composition of Strong vs. Weak Binders

The amino acid composition differed substantially between the top 10 and bottom 10 peptides (Figure 5, Table 3). Strong binders contained 38% charged and 40% polar residues, whereas weak binders were 75% hydrophobic. Cysteine and glutamate showed the largest enrichment (∼9-fold each, comparing top vs. bottom deciles). Arginine and threonine were also enriched (∼7-fold). Conversely, valine, glycine, alanine, phenylalanine, and tryptophan were depleted 3- to 7-fold in strong binders.

**Table 3.** Residue composition by physicochemical class.

| Class | Top 10 Binders | Bottom 10 Binders | Difference |
| --- | --- | --- | --- |
| Charged (D, E, K, R, H) | 15 (37.5%) | 4 (10.0%) | +27.5% |
| Polar (S, T, N, Q, C, Y) | 16 (40.0%) | 6 (15.0%) | +25.0% |
| Hydrophobic (A, V, L, I, P, F, W, M, G) | 9 (22.5%) | 30 (75.0%) | -52.5% |

**Figure 5.**
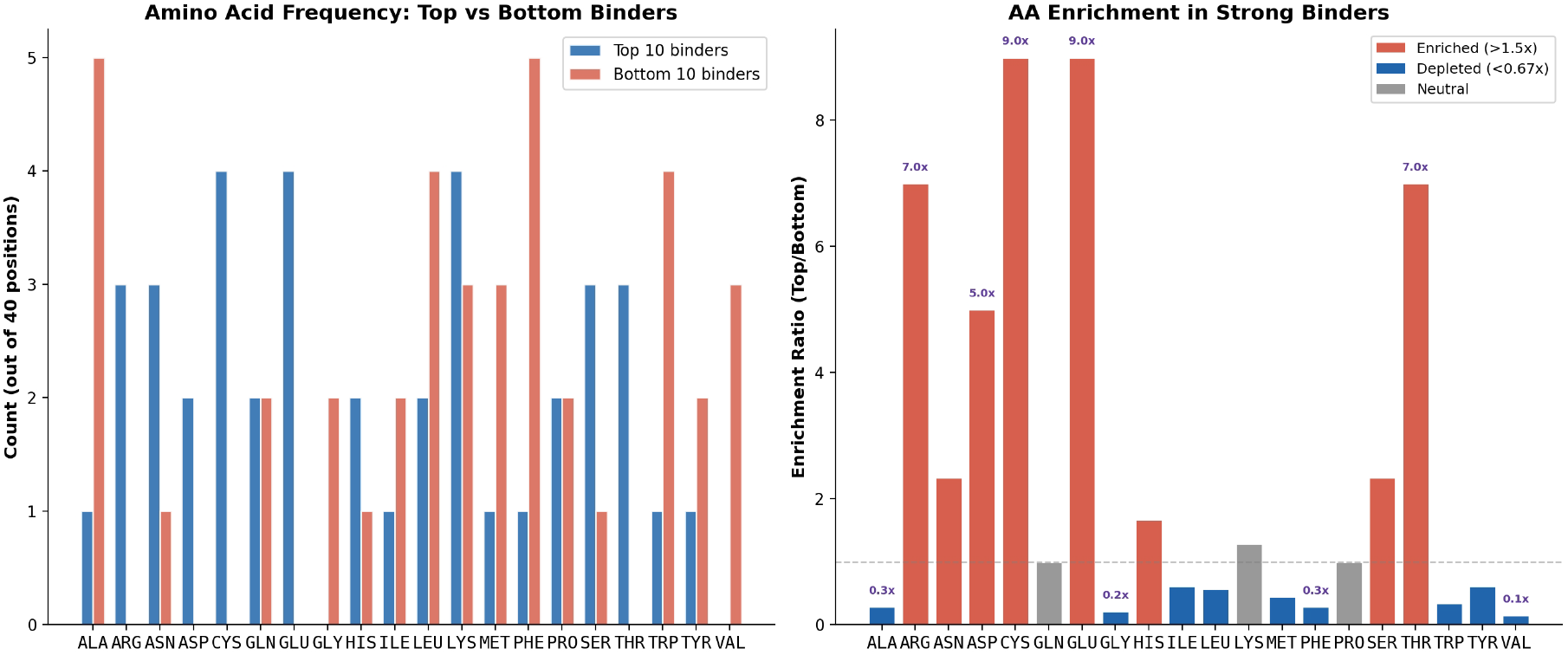
Amino acid analysis. (A) Frequency comparison between top-10 and bottom-10 peptides. (B) Enrichment ratios (top/bottom). Red: enriched >1.5×; blue: depleted <0.67×.

This pattern is the inverse of what Wang et al. reported for tetrapeptide aggregation, where aromatic and hydrophobic residues drive self-assembly and charged residues suppress it [8]. The residues that appear to promote IL-17A binding in our simulations are the same ones that keep tetrapeptides soluble.

### 3.5 Structural Dynamics

Peptide backbone RMSD averaged 0.30 ± 0.06 nm (range 0.16-0.47 nm) and showed little correlation with binding score (r = −0.12). Some peptides with high scores maintained relatively rigid bound conformations (LYS-THR-CYS-ASP: 0.25 nm), while others were more flexible (CYS-LYS-LYS-SER: 0.43 nm). IL-17A backbone RMSD averaged 2.12 ± 0.69 nm, reflecting the inherent flexibility of the cytokine’s loop-rich architecture [12,13]. Peptide binding did not appear to induce large-scale structural changes in IL-17A (Figure 6).

**Figure 6.**
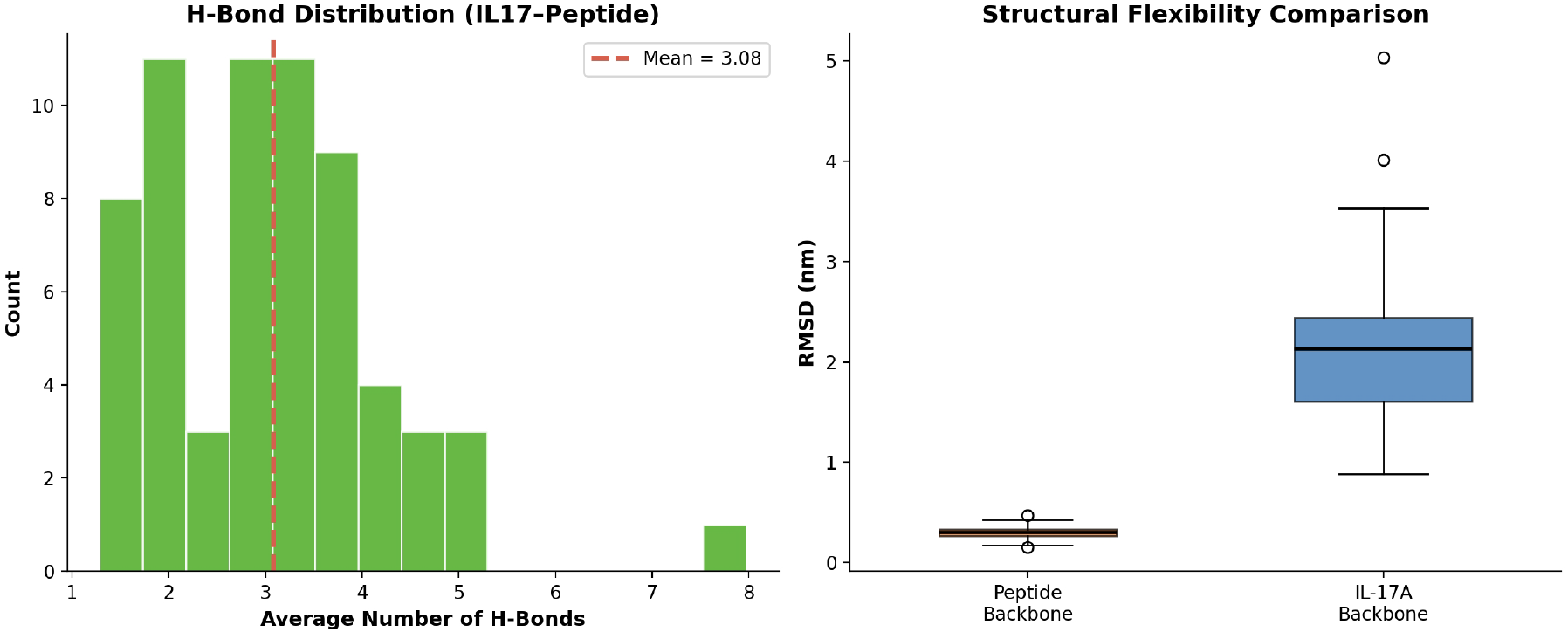
Structural metrics. (A) Distribution of average H-bond counts (mean = 3.08 ± 1.19). (B) Boxplot comparing peptide backbone RMSD and IL-17A backbone RMSD across all 64 simulations.

### 3.6 Correlation Structure of Binding Metrics

The Pearson correlation matrix among seven computed metrics (Figure 7) shows that the composite binding score is most closely related to total interaction energy (r = 0.78). Total energy and Coul-SR are nearly collinear (r = 0.93), consistent with the dominant electrostatic contribution. H-bond count correlates with Coul-SR (r = 0.72). Peptide RMSD is essentially uncorrelated with all other metrics. The modest correlation between LJ-SR and Coul-SR (r = 0.31) indicates that van der Waals and electrostatic terms capture partially independent aspects of the interaction [11,14].

**Figure 7.**
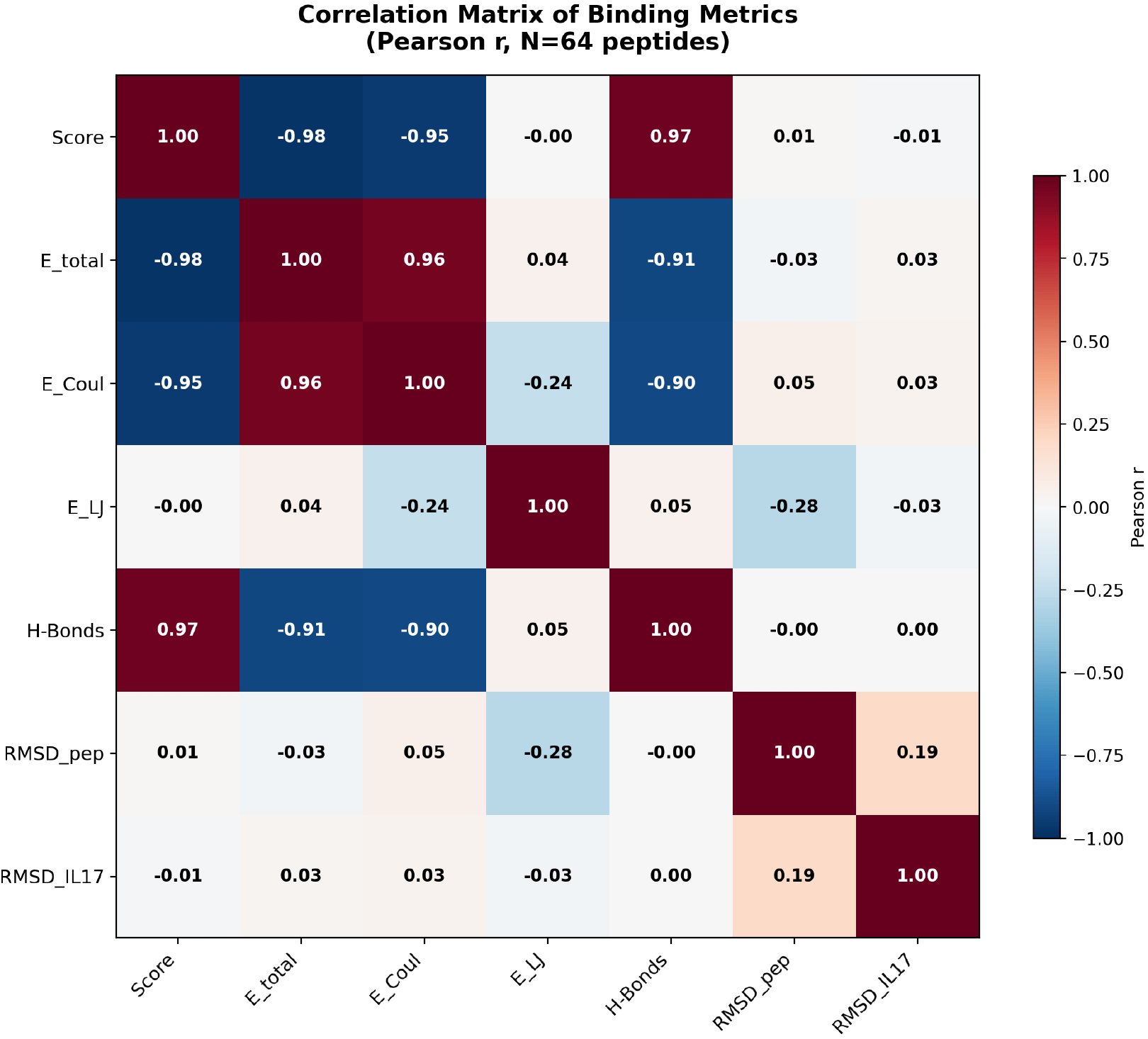
Pearson correlation matrix of seven binding-related metrics (N = 64).

## 4. Discussion

### 4.1 Features of LYS-THR-CYS-ASP

LYS-THR-CYS-ASP scored highest among the 64 peptides tested. Figure 8 summarizes its residue-level interactions with IL-17A. The four residue positions provide complementary interactions with the binding surface:

**Figure 8.**
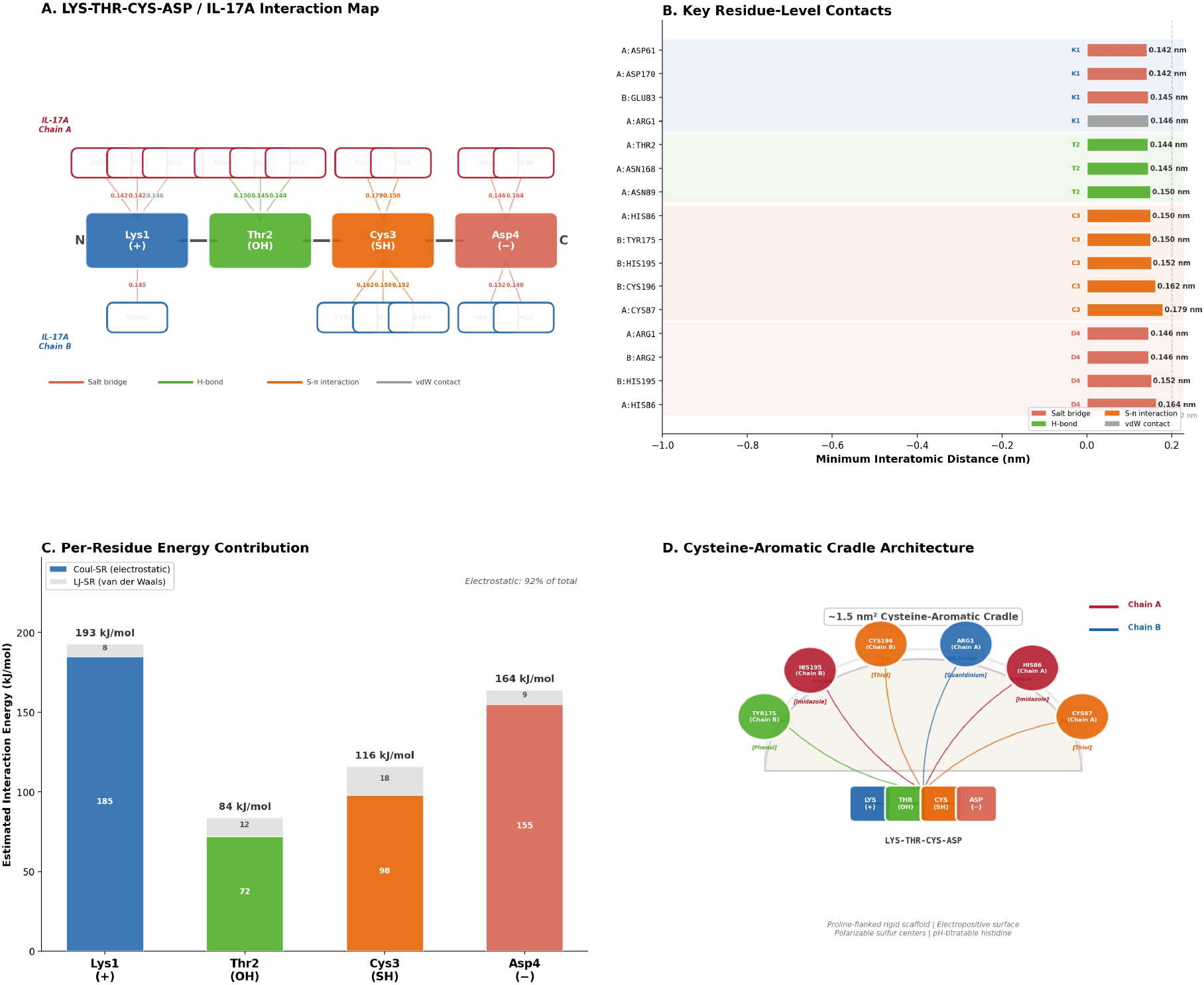
Interaction structure of LYS-THR-CYS-ASP with IL-17A. (A) 2D interaction map showing the tetrapeptide and its IL-17A contacts, colored by interaction type. (B) Minimum interatomic distances for each annotated contact. (C) Estimated per-residue energy contributions (Coul-SR + LJ-SR). (D) Schematic of the cysteine-aromatic cradle surrounding the bound tetrapeptide.

Lys1 forms salt bridges with acidic IL-17A residues (ASP61: 0.142 nm; ASP170: 0.142 nm; GLU83: 0.145 nm). The flexible lysine side chain permits conformational sampling while maintaining electrostatic contact.

Thr2 contributes hydrogen bonds to ASN89 (0.150 nm), ASN168 (0.145 nm), and THR2 (0.144 nm). As the smallest chiral H-bond-capable side chain, it bridges the peptide backbone to the protein surface with minimal entropic cost.

Cys3 positions its polarizable sulfur within the cysteine-aromatic cradle (CYS87: 0.179 nm; CYS196: 0.162 nm), where it can engage S–H···π interactions with HIS86, TYR175, and HIS195.

Asp4 anchors the C-terminus to basic hotspot residues (ARG1: 0.146 nm; HIS86: 0.164 nm; HIS195: 0.152 nm) through charge–charge interactions.

The net-neutral charge distribution (Lys^+^–Thr–Cys–Asp^−^) creates a dipole aligned with the electrostatic surface of the cradle. This N-to-C arrangement of positive– polar–thiol–negative charges may represent a useful pharmacophore pattern for targeting this site.

### 4.2 Sequence Preferences and Comparison with Aggregation

A practical observation from this screen is that the sequence features associated with stronger IL-17A binding in our simulations are approximately the opposite of those that promote tetrapeptide aggregation [8,9]. Aggregation is driven by aromatic and hydrophobic residues; IL-17A binding in our data favors charged and polar residues. If confirmed experimentally, this anti-correlation is convenient from a drug-development perspective, because candidate peptides would tend to have good aqueous solubility. Simple guidelines for designing IL-17A-targeting tetrapeptides based on our data include: (i) include at least one acidic residue (Asp or Glu); (ii) include at least one H-bond-capable residue (Cys, Thr, Ser, or Asn); (iii) consider cysteine at positions 2 or 3 to engage the cradle; and (iv) valine, alanine, phenylalanine, tryptophan, and glycine were rarely found among strong binders in this set.

### 4.3 The Cysteine-Aromatic Cradle

The IL-17A surface patch contacted by strong binders — centered on CYS87, HIS86, CYS196, HIS195, and TYR175 — is a chemically distinctive site. It is smaller and more localized than the binding footprint of the 15-residue HAP antagonist characterized by Liu et al. [5], and may represent a minimal recognition element within that larger interface. The cradle has several features that would be attractive for inhibitor design if binding can be confirmed experimentally: proline-flanked loops provide a relatively rigid scaffold; the chemical diversity in a compact area (salt-bridge partners, π systems, polarizable sulfur) enables multiple non-covalent interaction modes; and the solvent-accessible cysteine thiols raise the possibility of covalent inhibitor approaches, though this would require careful selectivity profiling.

### 4.4 Limitations

#### (1) Library size

64 tetrapeptides represent 0.04% of the possible 160,000 sequences. The results should be understood as screening a small, diverse sample, not a comprehensive search.

#### (2) Force field accuracy

FF19SB uses fixed atomic charges and cannot capture electronic polarization effects at the sulfur- and aromatic-rich interface. Polarizable force fields or QM/MM calculations would be needed for more accurate electrostatics.

#### (3) Sampling

200 ns per complex may be insufficient to fully sample binding and unbinding events. The trajectories reflect local exploration around the initial placement rather than global binding/unbinding equilibrium. Free energy methods (umbrella sampling, alchemical FEP) would be needed for quantitative binding affinity predictions.

#### (4) No entropy estimate

Our scoring function uses interaction energies and geometric metrics but does not account for binding entropy (conformational, translational/rotational, or solvent). MM/PBSA or MM/GBSA with entropy corrections would provide a more complete thermodynamic picture.

#### (5) Experimental validation needed

All results are computational predictions. The IL-17A structure is an AlphaFold3 model and should be compared with available crystal structures (PDB 4HR9, 4HSA) [12,13]. Binding must be tested by SPR, ITC, or cell-based assays before any candidate can be considered validated.

## 5. Conclusion

We used classical all-atom MD simulations to screen 64 tetrapeptides for binding to IL-17A. Electrostatic interactions (Coul-SR) accounted for roughly 72% of the total interaction energy across the dataset. Stronger binders were enriched in charged and polar residues (Asp, Glu, Cys, Arg, Thr), while weaker binders were dominated by hydrophobic residues. The IL-17A surface contacted by strong binders forms a cysteine-aromatic cradle (A:CYS87, B:CYS196, A:HIS86, B:TYR175, B:HIS195) that is electropositive and polarizable. The tetrapeptide LYS-THR-CYS-ASP achieved the highest composite score in our screen and represents a candidate for experimental testing. The sequence preferences observed here may be useful for designing focused peptide libraries for future computational or experimental screening campaigns.

## Acknowledgments

Simulations were run on the institutional HPC cluster (SLURM, amd_256 partition, AMD EPYC 64-core nodes) using GROMACS 2022.2 with Intel oneAPI 2022.1. We thank the HPC support team for infrastructure maintenance.

